# Diverse lineages of *Candida albicans* live on old oaks

**DOI:** 10.1101/341032

**Authors:** Douda Bensasson, Jo Dicks, John M. Ludwig, Christopher J. Bond, Adam Elliston, Ian N. Roberts, Stephen A. James

## Abstract

The human pathogen, *Candida albicans*, is considered an obligate commensal of animals, yet it is occasionally isolated from trees, shrubs and grass. We generated deep genome sequence data for three strains of *C. albicans* that we isolated from oak trees in an ancient wood-pasture, and compared these to the genomes of the type strain and 21 other clinical strains. *C. albicans* strains from oak are similar to clinical *C. albicans* in that they are predominantly diploid and can become naturally homozygous at the mating locus through whole-chromosome loss of heterozygosity (LOH). LOH regions in all genomes arose recently suggesting that LOH mutations usually occur transiently in *C. albicans* populations. Oak strains differed from clinical strains in showing less LOH, and higher levels of heterozygosity genome-wide. Using phylogenomic analyses, *in silico* chromosome painting, and comparisons with thousands more *C. albicans* strains at seven loci, we show that each oak strain is more closely related to strains from humans and other animals than to strains from other oaks. Therefore, the isolation of *C. albicans* from oak is not easily explained as contamination from a single animal source. The high heterozygosity of oak strains could arise as a result of reduced mitotic recombination in asexual lineages, recent parasexual reproduction or because of natural selection. Regardless of mechanism, the diversity of *C. albicans* on oaks implies that they have lived in this environment long enough for genetic differences from clinical strains to arise.

## Introduction

*Candida albicans* is the most common yeast pathogen of humans (Barnett, 2008). Yet, it is also a commensal in most humans and inhabits a broad range of warm-blooded animals (Barnett, 2008). Unlike other *Candida* species, *C. albicans* is only rarely isolated from plants, soil or other environmental substrates (Barnett, 2008; Lachance *et al.*, 2011) and is generally considered an “obligate commensal” (Hall and Noverr, 2017). There were a couple of early discoveries of *C. albicans* on gorse flowers and myrtle leaves on a hillside grazed by sheep and goats in Portugal (van Uden *et al.*, 1956), and on grass in a sheep pasture in New Zealand (Di Menna, 1958). More recently it was isolated from a beetle and an African tulip tree in the Cook Islands (Lachance *et al.*, 2011), and we isolated *C. albicans* from oak trees in an ancient wood-pasture in the United Kingdom (Robinson *et al.*, 2016).

Many species of yeast live on trees, including other *Candida* species (Maganti *et al.*, 2011; Charron *et al.*, 2014; Sylvester *et al.*, 2015) and woodlands represent the ancestral habitat for *Saccharomyces* species (Eberlein *et al.*, 2015). Forests may also be an ancestral habitat and reservoir for fungal pathogens that infect humans such as *Cryptococcus neoformans* and *Cryptococcus gattii* (May *et al.*, 2016; Gerstein and Nielsen, 2017). The isolation of *C. albicans* from plants (van Uden *et al.*, 1956; Di Menna, 1958; Lachance *et al.*, 2011; Robinson *et al.*, 2016) raises the question of whether *C. albicans* is truly an obligate commensal of warm-blooded animals. Lab experiments show that *C. albicans* can grow and mate at room temperature (Magee and Magee, 2000; Hull *et al.*, 2000) and it retains an intact aquaporin gene whose only known phenotype is freeze tolerance (Tanghe *et al.*, 2005). It is therefore possible that *C. albicans* populations could survive away from warm-blooded animals.

Here we generated genome sequences for three strains of *C. albicans* from oak bark in the New Forest in the U.K. (Robinson *et al.*, 2016) and compared them to the genomes of a well-studied panel of clinical strains (Wu *et al.*, 2007; Sahni *et al.*, 2009; Muzzey *et al.*, 2013; Hirakawa *et al.*, 2015; Wang *et al.*, 2018). The three oak strains are genetically diverged from one another, and are more similar to clinical strains than they are to each other. However, genomes from oaks do differ from those of clinical strains in that they show higher levels of genome-wide heterozygosity. The genetic diversity of *C. albicans* in this oak woodland cannot easily be explained as contamination from a human source, and suggests that *C. albicans* can live in the oak environment for extended periods of time.

## Results

### Phenotypically diverse strains of *C. albicans* from old oaks

We recently discovered *C. albicans* living on the bark of oak trees in a wood-pasture in the New Forest, and the three strains we isolated are available from the U.K. National Collection of Yeast Cultures (NCYC 4144, NCYC 4145, NCYC 4146; Table 1, Robinson *et al.*, 2016). For several reasons, the data from Robinson *et al.* (2016) suggest that these three oak strains represent independent isolates and not contaminants from a single human or animal source. Bark samples were collected into tubes using sterile technique from heights of at least 1.5 meters above the ground, thus reducing the chances of direct contamination from animal manure. Negative controls generated in the field at the time of collection gave rise to no colonies after enrichment culturing, and subsequent DNA extraction and PCR amplification from these control plates were all also blank (Robinson *et al.*, 2016). The three trees harboring *C. albicans* were between 73 and 150 meters apart, therefore any migration from animal manure or from humans to tree bark would have to occur in three separate events. Finally, the trees harboring *C. albicans* had larger trunk girths and therefore were older than most of the 112 uncoppiced trees sampled across Europe (data from Robinson *et al.* 2016, Wilcoxon test, *P* = 0.009) or in the New Forest (Table 1; Wilcoxon test, *P* = 0.04). There is no reason to expect a greater level of contamination from humans on old trees unless *C. albicans* are able to live on oaks for many years.

**Table 1:**
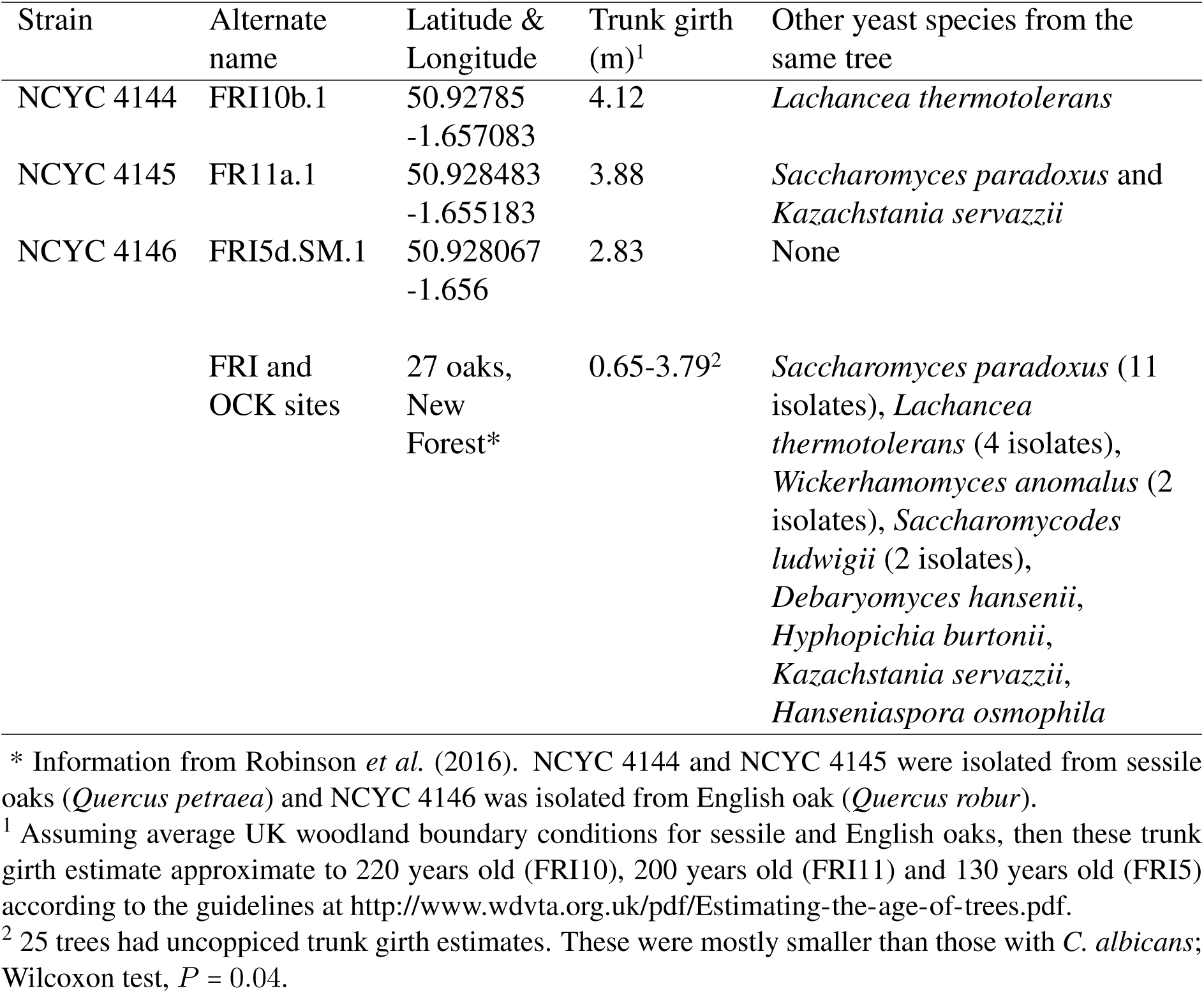
Three *C. albicans* isolates from English and sessile oaks in the New Forest in the United Kingdom*.

*C. albicans* that were isolated from plants in the past were pathogenic in mammals, but did not differ from each other phenotypically (van Uden *et al.*, 1956; Di Menna, 1958). In order to test whether this is also true for the *C. albicans* strains isolated from oak, we tested the growth of oak strains under the standard conditions described in (Kurtzman *et al.*, 2011). All three oak strains were able to grow at elevated temperatures (37-42°C), suggesting that they would be able to survive in a mammalian host. The oak strains also produced well-formed, irregularly branched pseudohyphae when grown on either corn meal agar or potato dextrose agar, and grew in 60% glucose, showing that they were highly osmotolerant. The phenotypes of the three oak strains differed from each other in that NCYC 4144 and NCYC 4146 displayed salt tolerance by growing in the presence of 10% NaCl, but NCYC 4145 did not. The results of further growth tests on agar, in broth and under other conditions were as previously described for *C. albicans* (Lachance *et al.*, 2011) with the following exceptions: (i) one oak strain (NCYC 4144) formed pseudohyphae in YM broth and did not grow on soluble starch; (ii) another oak strain (NCYC 4145) was unable to grow on galactose.

In addition, the three *C. albicans* strains from oak must be able to survive the cool temperatures and other characteristics of the oak environment because they lived there until they were isolated from bark. They also survived enrichment, storage and culturing conditions, which include growth at room temperature, 30°C, in a liquid medium containing chloramphenicol (1 mg/l) and 7.6% ethanol, on selective plates with a sole carbon source of methyl-*α*-D-glucopyranoside (Sniegowski *et al.*, 2002), storage at 4°C and as 15% glycerol stocks at −80°C (Robinson *et al.*, 2016).

### *C. albicans* from oak are mostly diploid

Clinical strains of *C. albicans* are predominantly diploid (Hickman *et al.*, 2013; Hirakawa *et al.*, 2015), yet aneuploidy is often observed and may be an important mechanism for adaptation (Li *et al.*, 2015; Todd *et al.*, 2017). We compared oak and clinical strain ploidy by comparing the short-read genome sequence data generated here to published data for laboratory and clinical strains. More specifically, we applied a standard base calling approach for estimating ploidy from sequences to data for oak strains, the laboratory reference strain (SC5314), a related mutant (1AA) (Muzzey *et al.*, 2013), and 20 clinical strains (Hirakawa *et al.*, 2015). In addition, we generated short-read genome data for the clinical type strain of *C. albicans* (NCYC 597) for a direct comparison between oak strains and a clinical strain from the same sequencing batch.

The base calling approach we used (the B allele approach; Teo *et al.*, 2012; Yoshida *et al.*, 2013; Zhu *et al.*, 2016) estimates the minimum ploidy of a yeast strain by examining the base calls of short read genome data mapped to a reference genome. In a haploid genome or a homozygous diploid, base calls at the single nucleotide polymorphic (SNP) sites relative to the reference genome will all differ from the reference genome, so the proportion of base calls that differ from the reference will be approximately equal to 1 (e.g. chromosome 5 of the oak strain NCYC4144 in Figure 1). In a diploid, the proportion of base calls differing from the reference will be approximately equal to 1 at homozygous SNP sites or 0.5 at heterozygous SNP sites (e.g. the oak strain NCYC 4146 in Figure 1). In triploids, the proportion of calls differing from the reference will be 0.66 and 0.33 at heterozygous sites (e.g. the type strain in Figure 1) and so on. It is also possible to detect aneuploidy by comparing read depth between chromosomes. However, we use a base calling approach because read depth approaches are sensitive to biases when genomes are fragmented enzymatically as they were in this study (see Methods; Marine *et al.*, 2011; Quail *et al.*, 2012; Teo *et al.*, 2012).

**Figure 1:**
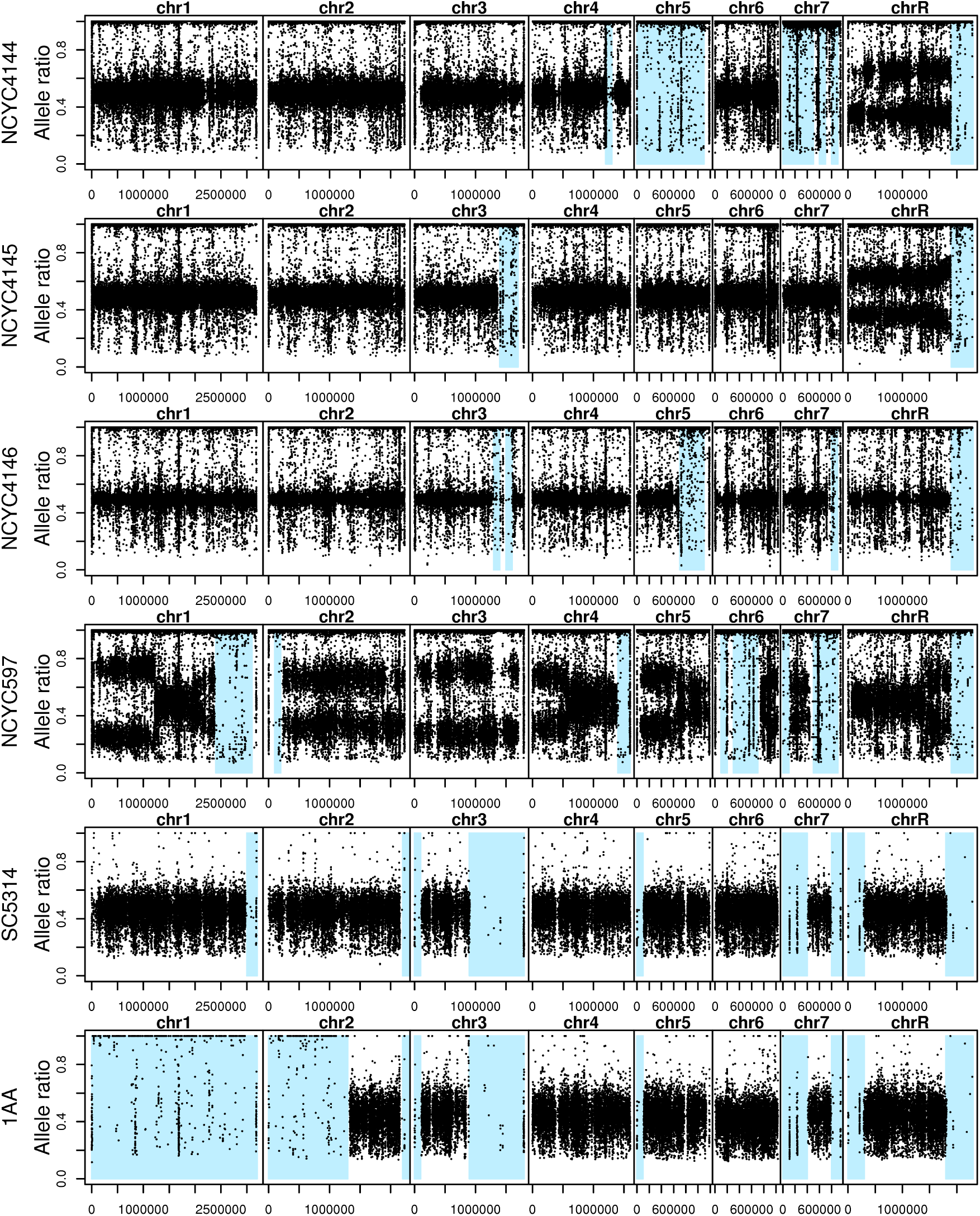
*C. albicans* from oak are mostly diploid, whereas the *C. albicans* type strain is mostly triploid. The proportion of base calls differing from the reference strain (allele ratios) are mostly 1.0 or 0.5 for oak strains (NCYC 4144-6) suggesting for diploidy, whereas allele ratios are mostly 0.33, 0.66 and 1.0 for the type strain (NCYC 597) suggesting triploidy. As expected, SC5314 differs from the SC5314 A22 reference at heterozygous sites, and the laboratory mutant (1AA) is homozygous on chromosome 1. Regions that recently homozygosed are shaded light blue. The points that occur in these loss of heterozygosity (LOH) regions often correspond to the locations of known repeats where short reads are probably mismapped and repeat regions were mostly filtered from final estimates of heterozygosity in Table 2. The oak strain with *a/a* at its mating locus (NCYC 4144) arrived at this state by loss of heterozygosity for the whole of chromosome 5.

By applying the base calling approach to the laboratory and clinical strains analyzed by Muzzey *et al.* (2013) and Hirakawa *et al.* (2015), we confirm their conclusion that these strains are all predominantly diploid (Figure S1). However, this approach does not allow determination of ploidy state for whole chromosomes that have recently become homozygous. We were able to detect six out of the seven cases of trisomy identified by Hirakawa *et al.* (2015), but missed tetrasomy for chromosome 5 in strain P75010 which was was fully homozygous (File S1). Therefore we could miss aneuploidy for homozygous chromosomes, but there are few cases of this for the oak strains (only chromosomes 5 and 7 of strain NCYC 4144 from oak, Figure 1).

In contrast to the clinical strains studied by Hirakawa *et al.* (2015), the type strain of *C. albicans* is predominantly triploid (Figure 1). Therefore, we could have detected ploidy variation in the oak strains had it been present. Indeed, our data suggest that the type strain has probably undergone large scale chromosomal rearrangements because most of its chromosomes show a mixture of ploidy states along their sequence (chromosomes 1, 4, 5, 6 and R, Figure 1).

Our analysis of base calls in oak strains suggests that all three strains are predominantly diploid (Figure 1). NCYC 4146 appears to be euploid, but NCYC 4144 and NCYC 4145 show evidence of trisomy on chromosome R. The type strain and one clinical strain (P60002) also have three copies of the right arm of chromosome R (Hirakawa *et al.*, 2015; Figures 1 and S1). The right arm of chromosome R may therefore exist in three copies with appreciable frequency in natural strains of *C. albicans*. Although trisomy of chromosome R can result in slow growth in the laboratory (Hickman *et al.*, 2015), it has been associated with resistance to triazoles (Li *et al.*, 2015).

### Recent loss of heterozygosity in oak and clinical strains

The base calling approach we used to determine overall ploidy also shows that *C. albicans* are mostly highly heterozygous diploids with interspersed regions that have recently undergone loss of heterozygosity (LOH; Figures 1 and S1). LOH events often occur in *C. albicans* genomes by mitotic recombination or loss of whole chromosomes, they affect mating type and could have important effects on other phenotypes (Bougnoux *et al.*, 2006, 2008; Forche *et al.*, 2011; Hirakawa *et al.*, 2015; Ford *et al.*, 2015). We therefore tested whether oak strains are similar to clinical strains in showing evidence of LOH in their genomes.

Using our methods, we detected multiple LOH events in every oak strain in addition to the previously known LOH events for clinical strains reported by (Hirakawa *et al.*, 2015) (Figure S1), and to the artificially induced whole-chromosome LOH of chromosome 1 for the mutant strain 1AA (Figure 1). Read depth in regions with low heterozygosity (Figures 1 and S1) is similar to that in the rest of the genome (Figure S2), therefore these regions probably do not represent deletions or the loss of a chromosome. Consistent with the proposal that aneuploidy persists for less time in *C. albicans* populations than LOH regions (Ford *et al.*, 2015), we observe a greater number of LOH regions than aneuploidy events for every clinical or oak strain (Figures 1 and S1).

For both oak and clinical strains, the length of LOH regions vary from short chromosomal segments to whole chromosomes (Figures 1 and S1). Indeed, one oak strain (NCYC4144) is homozygous across the whole of chromosome 5 on which the MTL locus is situated. Analysis of whole-genome data, and confirmation by independent PCR and sequencing shows that this strain is homozygous for the *a* allele at the *C. albicans* mating locus (*a/a*) and therefore could potentially mate with strains that are homozygous for the opposite mating type (*α/α*). Whole-chromosome homozygosis is not an unusual mechanism by which natural strains of *C. albicans* become homozygous at the MTL locus (Hirakawa *et al.*, 2015). Two out of the 10 naturally occurring MTL-homozygous clinical strains included in this study are also homozygous across the whole of chromosome 5 (Figure S1; Sahni *et al.*, 2009; Hirakawa *et al.*, 2015).

After an LOH event occurs, the resulting homozygous region gradually accumulates new mutations, therefore levels of heterozygosity within an LOH region are an indication of the age of an LOH event. If LOH is an ongoing mechanism in *C. albicans* genome evolution (Bougnoux *et al.*, 2008), then we would expect *C. albicans* genomes to vary continuously in their levels of heterozygosity. However, for both oak and clinical strains, most of the genome shows high heterozygosity (over 0.4% of nucleotide sites are heterozygous), or low heterozygosity (below 0.1%), but relatively few regions have intermediate heterozygosity levels (Figures 2 and S3). This bimodal distribution (Figures 2 and S3) therefore implies that most LOH regions in *C. albicans* genomes arose recently. Even if LOH is an important mechanism for rapid adaptation to environmental stress (Forche *et al.*, 2011; Gerstein *et al.*, 2014), its recent origins in all the strains studied here suggest that most LOH regions only exist transiently in populations.

**Figure 2:**
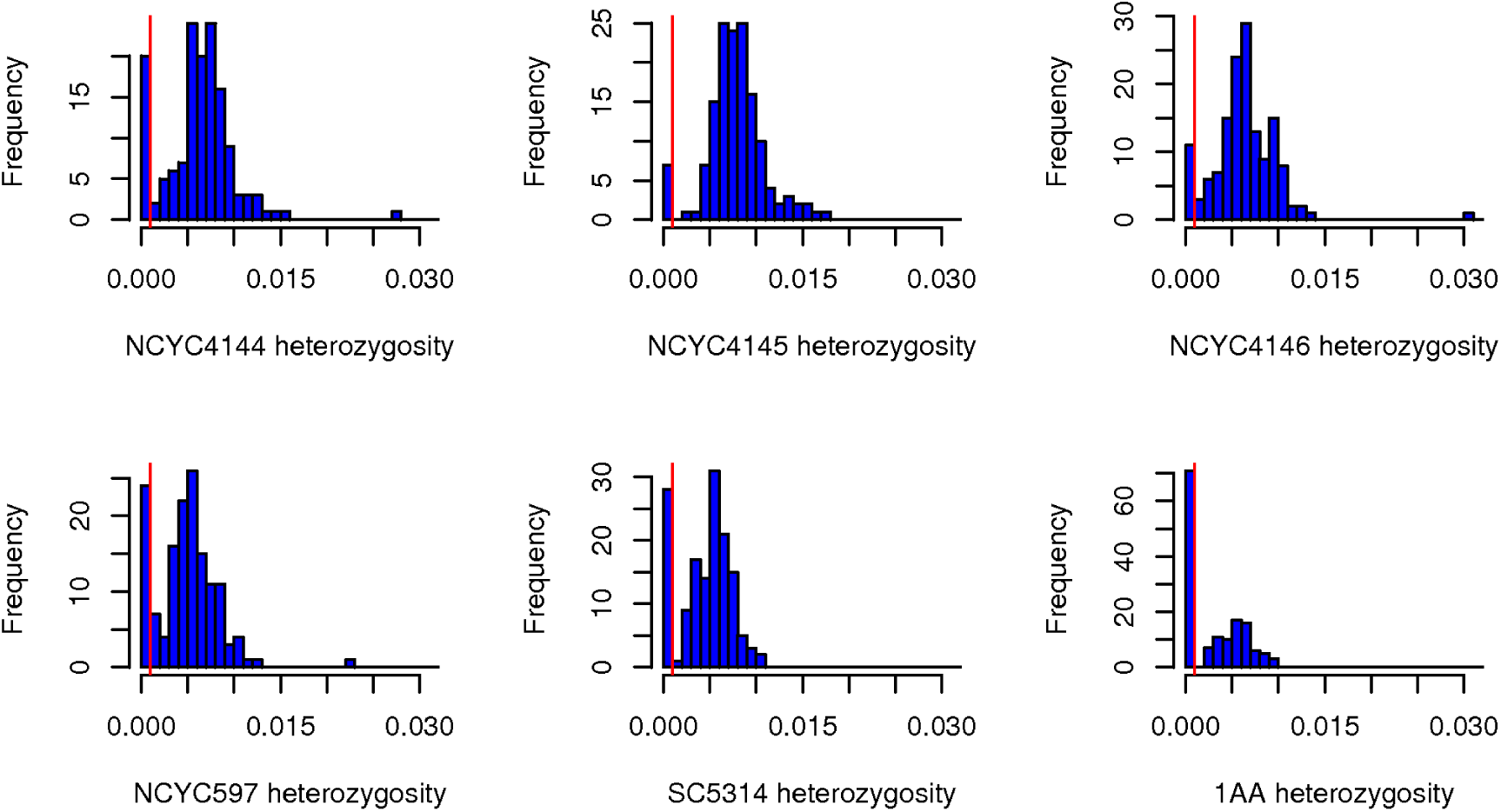
Levels of heterozygosity in 100 kb regions are either high or low for oak and clinical strains. The proportion of heterozygous sites was estimated in 100 kb non-overlapping windows across the genome of each strain. Results are shown here for oak strains (NCYC 4144, NCYC 4145, and NCYC 4146), the type strain (NCYC 597), the wild type version of the laboratory strain used to generate the reference genome for *C. albicans* (SC5314) and a mutant that was made homozygous for chromosome 1 in the laboratory (1AA, Legrand *et al.*, 2008). Results for 20 more clinical strains are shown in Figure S3. For all strains we see two modes; heterozygosity is either low (below the red line at 0.1%), or high (with a mean above 0.4%). Regions with fewer than 0.1% heterozygous sites in a 100 kb window were classed as LOH regions and are shown in blue in Figure 1.

### High heterozygosity on oaks

High levels of genome-wide heterozygosity can be an indicator of prolonged asexuality (Birky, 1996; Halkett *et al.*, 2005). Therefore clonal divergence could explain why levels of heterozygosity are high in clinical *C. albicans* strains (Bougnoux *et al.*, 2008). This is because after loss of sex and in the absence of mitotic recombination, the haplotypes within a lineage will diverge as they accumulate mutations. In contrast, in sexual species, meiotic recombination can lead to increased similarity between alleles (Birky, 1996; Halkett *et al.*, 2005). Because differences in levels of heterozygosity could reveal differences in the frequency of asexual or parasexual life cycles, we compared levels of heterozygosity between oak and clinical strains. For every *C. albicans* strain, we obtained high quality sequence for approximately 14 million sites in the genome and estimated the proportion of these sites that were heterozygous (Tables 2 and S1).

**Table 2:** *C. albicans* from oak show higher heterozygosity than clinical strains.

| Strain | MTL | MLST<br>Clade<br>(FP<sup>a</sup>) | Heterozy-<br>gosity<sup>b</sup> | LOH<br>length<br>(Mbp)<sup>c</sup> | Filtered<br>length<br>(Mbp)<sup>d</sup> | Filtered<br>heterozy-<br>gosity<sup>e</sup> | Heterozy-<br>gosity in<br>950 kb<sup>f</sup> |
| --- | --- | --- | --- | --- | --- | --- | --- |
| NCYC 4145 | *a/α* | ? | 0.0077 | 0.7 | 11.4 | 0.0078 | 0.0075 |
| NCYC 4146 | *a/α* | 4 (SA) | 0.0062 | 1.1 | 11.1 | 0.0066 | 0.0068 |
| NCYC 4144 | *a/a* | ? | 0.0061 | 2.3 | 9.7 | 0.0068 | 0.0074 |
| **3 oak**<br>**strains** |  | Mean: | **0.0066** | **1.4** |  | **0.0070** | **0.0072** |
| P34048 | *a/α* | 3 (III) | 0.0060 | 1.0 | 10.3 | 0.0064 | 0.0070 |
| P75016 | *a/α* | 4 (SA) | 0.0059 | 0.9 | 10.5 | 0.0064 | 0.0066 |
| P78042 | *α/α* | 3 (III) | 0.0057 | 1.3 | 10.1 | 0.0061 | 0.0065 |
| GC75 | *α/α* | 4 (SA) | 0.0057 | 1.6 | 9.4 | 0.0067 | 0.0067 |
| P78048 | *α/α* | 1 (I) | 0.0054 | 2.1 | 9.5 | 0.0061 | 0.0054 |
| P57055 | *a/α* | 3 (III) | 0.0052 | 3.0 | 8.9 | 0.0065 | 0.0068 |
| P37037 | *a/α* | 1 (I) | 0.0048 | 3.3 | 8.3 | 0.0061 | 0.0058 |
| NCYC597 | *a/α* | ? | 0.0048 | 2.4 | 9.3 | 0.0054 | 0.0061 |
| P37005 | *a/a* | 1 (I) | 0.0047 | 3.3 | 8.4 | 0.0059 | 0.0057 |
| P57072 | *α/α* | 2 (II) | 0.0047 | 2.7 | 8.4 | 0.0057 | 0.0058 |
| P75010 | *α/α* | 11 (E) | 0.0046 | 4.7 | 6.7 | 0.0067 | 0.0068 |
| SC5314 | *a/α* | 1 (I) | 0.0046 | 2.8 | 10.0 | 0.0055 | 0.0053 |
| P76067 | *a/α* | 2 (II) | 0.0046 | 3.9 | 8.6 | 0.0061 | 0.0055 |
| P37039 | *a/α* | 1 (I) | 0.0045 | 3.9 | 8.1 | 0.0060 | 0.0057 |
| P75063 | *a/α* | 4 (SA) | 0.0045 | 3.6 | 8.1 | 0.0059 | 0.0067 |
| L26 | *a/a* | 1 (I) | 0.0044 | 3.7 | 9.2 | 0.0058 | 0.0056 |
| 19F | *α/α* | 1 (I) | 0.0042 | 4.3 | 8.1 | 0.0059 | 0.0057 |
| P76055 | *a/α* | 2 (II) | 0.0042 | 3.5 | 8.3 | 0.0052 | 0.0052 |
| P60002 | *a/a* | 8 (SA) | 0.0041 | 4.4 | 7.7 | 0.0056 | 0.0070 |
| 12C | *a/a* | 1 (I) | 0.0041 | 4.7 | 7.7 | 0.0061 | 0.0057 |
| P87 | *a/a* | 4 (SA) | 0.0038 | 5.7 | 6.8 | 0.0062 | 0.0060 |
| P94015 | *a/a* | 6 (I) | 0.0035 | 6.5 | 6.8 | 0.0062 | 0.0068 |
| 1AA | *a/α* | 1 (I) | 0.0029 | 7.1 | 5.6 | 0.0056 | 0.0039 |
| **22 clinical**<br>**strains<sup>g</sup>** |  | Mean: | **0.0047** | **3.3** |  | **0.0060** | **0.0061** |

<sup>a</sup> Clade assignments are as summarized from past MLST and fingerprinting (FP) studies in Hirakawa *et al.* (2015). <sup>b</sup> Heterozygosity was estimated as the proportion of high quality sites (with phred-scaled quality over 40) where 20-80% of reads differed from the reference sequence. For all strains, this was estimated from approximately 14 Mbp of high quality sequence. <sup>c</sup> Length of sequence showing loss of heterozygosity (LOH). LOH was assumed where the proportion of heterozygous sites in a 100 kb window was lower than 0.001. <sup>d</sup> The length of genome sequence after excluding LOH regions, known repeats, putatively repetitive regions (positions with over double the mean genome-wide read depth) and centromeres. <sup>e</sup> The proportion of heterozygous sites after excluding LOH regions, repeats and centromeres. <sup>f</sup> the proportion of heterozygous sites in at 948,860 nucleotide sites with high quality, unreplicative, non-LOH sequence for all 25 oak and clinical strains. <sup>g</sup> Means for clinical strains exclude data for strain 1AA because this strain was derived from SC5314.

Levels of heterozygosity are higher in the 3 oak strains (0.61-0.77%) than they are for clinical strains (0.35-0.60%, Table 2; Wilcoxon test, *P* = 0.0009). In contrast, the clinical strain of *C. albicans* (NCYC 597) that we studied in the same sequencing batch as the oak strains showed a level of heterozygosity (0.48%) that was similar to other clinical strains, suggesting that high heterozygosity is not an artifact of the sequencing methods used in this study (Table 2). Furthermore, we excluded all sites with low quality sequence (with an expected error rate over 1 in 10,000). We generated more high quality sequence for all three oak strains (14,072,669 - 14,235,230 bp) compared to this control clinical strain (13,948,647 bp), and the oak strains did not differ from clinical strains in the amount of high quality sequence analyzed (14,184,615 - 14,259,261 bp; Wilcoxon test, *P* = 0.1; Table S1).

The high heterozygosity of oak strains compared with clinical strains could be caused by a difference in the amount of the genome that shows recent LOH. Even though the oak strain with the *a/a* mating type (NCYC4144) has undergone recent LOH for multiple chromosomes, oak strain genomes show less LOH than those of clinical strains (Wilcoxon test, *P* = 0.03; Table 2, Figures 1 and S1).

Levels of heterozygosity could differ in centromeres which evolve faster than other genomic regions in *C. albicans* (Padmanabhan *et al.*, 2008) and in other yeast species (Bensasson *et al.*, 2008). The number of heterozygous sites could also be overestimated in repetitive regions as a result of the mismapping of short reads to the reference genome. We therefore estimated levels of heterozygosity after filtering out centromeres, known repeats (using the reference genome annotation), and sites with over double the mean genome-wide read depth that could represent unannotated repeats. Even after excluding LOH regions, centromeres and repeats, levels of heterozygosity are higher for oak strains (mean 0.70%) than they are for clinical strains (mean 0.60%; Wilcoxon test, P = 0.003, Table 2). This difference in heterozygosity results from heterozygosity at thousands of sites across the genome (Table S1). For example, the strain NCYC 4145 is heterozygous at 0.78% of the 11.4 million sites we studied after filtering, and therefore has over 20,000 more heterozygous sites than expected for a clinical strain, and over 12,000 more heterozygous sites than expected for the most heterozygous clinical strains (0.67% for GC75 and P75010). Once more, this is not explained by a difference in sequence quality, because there is no correlation between the total length of high quality sequence generated and levels of filtered heterozygosity (Pearson’s correlation; *ρ* = −0.04; *P* = 0.8).

After filtering LOH regions, centromeres and repeats, heterozygosity was estimated from a larger component of the genome for oak (9.7-11.4 Mbp) compared to clinical strains (6.8- 10.5 Mbp). Could the longer component analyzed for oak strains include faster evolving regions that explain the higher heterozygosity seen for oak strains? To address this question, we identified 948,860 nucleotide sites that had not undergone LOH in any strain except 1AA, had high quality sequence for all 25 study strains, did not occur in centromeres and were not repetitive. We excluded the laboratory strain 1AA from all summary analyses of heterozygosity because this is an SC5314 derivative that had undergone artificially induced LOH. The resultant 948,860 nucleotide sites that were common to all 25 study strains were mostly on chromosomes 1, 2, 4 and 6. At these sites, oak strains were more heterozygous (mean 0.72%) than clinical strains (mean 0.61%; Wilcoxon test, P = 0.01; Table 2). This suggests that oak strains (especially NCYC 4144 and NCYC 4145; Table 2) show higher levels of heterozygosity than clinical strains throughout their genomes.

An unusually large proportion of the clinical strains are homozygous at the MTL locus (12 out of 22 strains). Could the low level of genome-wide heterozygosity in clinical strains result because this is a biased, unusually homozygous sample? After excluding LOH regions, centromeres and repeats from our analysis, we were unable to detect a difference in levels of genome-wide heterozygosity between MTL heterozygous clinical strains (mean 0.60%) and MTL homozygous clinical strains (mean 0.60%; Wilcoxon test, *P* = 0.8). Furthermore, the two *a/α* oak strains (NCYC 4145 and NCYC 4146) show higher levels of genome-wide heterozygosity (0.78% and 0.66%) than the ten *a/α* clinical strains (0.52%- 0.65%; Wilcoxon test, *P* = 0.03; Table 2). Therefore biased sampling of clinical strains for mating locus genotype does not explain the differences we see between clinical strains and oak strains.

In order to test whether the clinical genomes sampled here represent a biased sample of strains with respect to heterozygosity, we also compared levels of heterozygosity for the 22 clinical strains in our genome-wide sample to estimates for 1,391 clinical strains studied by Odds *et al.* (2007). The multilocus sequence typing (MLST) data of Odds *et al.* (2007) includes diploid sequence for a large global sample of *C. albicans* strains. After filtering out repeats, centromeres and LOH regions, our genome-wide estimates of heterozygosity (mean 0.60%) are similar to the heterozygosity estimates for these same clinical strains at MLST loci (mean 0.62%), and the same as the average level of heterozygosity at MLST loci estimated from 1,391 more clinical strains (mean 0.60%). Therefore the high levels of heterozygosity reported here are unlikely to be the result of the biased sampling of clinical strains for genome analysis.

### Oak strains are phylogenetically diverse

Most clinical strains of *C. albicans* belong to a small number of genetically diverged clades (Odds *et al.*, 2007). Strains belonging to the four most common clades (MLST clades 1-4 in Table 2 and Figure 3a) have a global distribution and live alongside each other in the same human populations (Bougnoux *et al.*, 2006; Odds *et al.*, 2007). Phylogenetic comparisons between oak and clinical strains can be used to determine whether oak strains form distinct populations that differ genetically from clinical strains; as they do in *S. cerevisiae* (Almeida *et al.*, 2015; Peter *et al.*, 2018). We therefore compared oak strains to clinical strains from seven of the most abundant *C. albicans* clades by whole-genome phylogenetic analysis.

As expected under clonality (Birky, 1996; Halkett *et al.*, 2005), phylogenies are congruent for most clinical strains whether we consider whole genomes, individual chromosomes or other genomic regions (Figure 3, S4, S5, and S6). We also painted the chromosomes of each clinical strain *in silico* according to the clade assignment of similar strains (Figures 3b and S7). This fine-scale analysis shows that clade assignments for clinical strains are consistent across almost all parts of the genomes we have studied (Figure S7). Our genome-wide analyses therefore support the conclusion that reproduction in clinical strains is predominantly asexual, or at least that there has been little recent gene flow between *C. albicans* clades in the case of the 18 clinical strains that Hirakawa *et al.* (2015) assigned to well-sampled clades.

**Figure 3:**
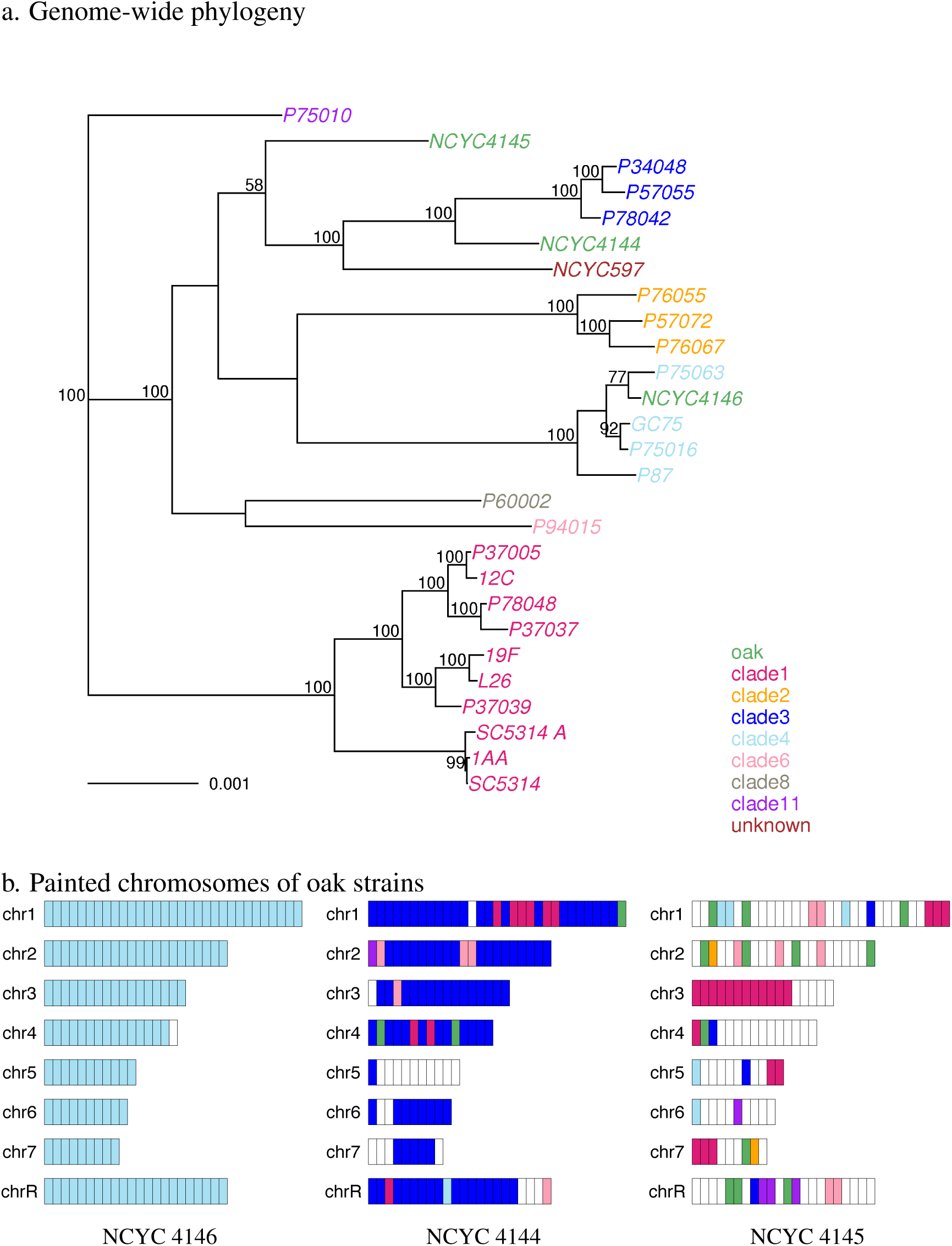
*C. albicans* from oak are more similar to clinical strains than to each other. Phylogenetic and pairwise sequence comparisons show that the oak strain NCYC 4146 is similar to clade 4 clinical strains (grey), NCYC 4144 is similar to clade 3 clinical strains, and NCYC 4145 is diverged from most sampled strains. a. Maximum likelihood phylogenetic analysis of whole genomes in a concatenated alignment shows that oak strains (green) are more closely related to clinical strains than they are too each other. b. Most parts of the genomes of oak strains are more similar to clinical strains than to other oak strains (green). The genome of each oak was coloured according to the clade assignment of the most similar strain for each 100 kb window in the genome. Regions are coloured white if a strain sequence is diverged from all the other oak or clinical strains that we sampled (the proportion of sites differing is over 0.066%).

All three oak strains are phylogenetically distinct from each other and more similar to clinical strains than they are to each other (Figure 3). Phylogenetic analyses of whole-genome data, separate chromosomes and subregions (Figures 3, S4, S5 and S6) all show that one strain from oak (NCYC 4146) belongs to MLST clade 4. This strain is also most similar to clade 4 strains throughout its genome (Figure 3b). In contrast, the other two strains from oak (NCYC4144 and NCYC4145) are diverged from each other and cannot be unambiguously assigned to any of the seven common *C. albicans* clades (Figures 3).

Moreover, these two oak strains (NCYC4144 and NCYC4145) showed different phylogenetic relationships with clinical strains in different parts of the genome (Figures 3b, S5, and S6). Analysis of one oak strain (NCYC4145) in short (100 kb) blocks across the whole genome suggests that it is diverged from the seven sampled clades (Figure 3b). Compared with other oak strains (Figure 3b) or with clinical strains from known clades (Figure S7a-d), NCYC4145 had more regions that were diverged from other sampled strains. In this, NCYC4145 is similar to clinical strains that are from clades only represented by a single strain (Figure S7e).

In contrast, the oak strain with the *a*/*a* mating type (NCYC4144) is mostly similar to clade 3, but shows some evidence of recent genetic admixture from other unidentified clades (Figure 3b). If a strain shows admixture between multiple clades, then heterozygosity at sites differing between the parental haplotypes will prevent recognition of their phylogenetic relationships. We therefore ran phylogenetic analyses in genomic regions where this strain is homozygous: chromosome 5, chromosome 7 and the right arm of chromosome R (Figure 1). These analyses suggest that NCYC4144 is more similar to other clades than it is to clade 3 strains in some homozygous parts of the genome (see example in Figure S6). However, NCYC4144 is also somewhat diverged from clade 3 and other known clades in all regions investigated (Figures 3b and S6) so it may represent a distinct clade with ancestral similarity to the other clades in this study. Without more extensive sampling of *C. albicans* strains and phased haplotype sequences, we cannot determine with certainty whether there has been genetic exchange between clades in the recent ancestry of this strain.

## Discussion

### High diversity of *C. albicans* from oaks

Phylogenetic analyses and fine-scale genome-wide DNA sequence comparisons show that all three strains from oaks from a single woodland site belong to distinct clades and therefore differ genetically as much as possible (Figures 3, S5, and S7). Consistent with genome-wide analyses, comparison of the MLST sequences of oak strains to over 3,000 sequences available for clinical strains (https://pubmlst.org/calbicans/) shows that each oak strain is similar at MLST loci to clinical strains from a different continent (U.K., U.S.A, China and South Korea; Supplemental Results). In this, oak strains are similar to *C. albicans* strains from wild and domestic animals. Three independent studies of *C. albicans* from Germany (Edelmann *et al.*, 2005), northwestern Europe (Jacobsen *et al.*, 2008) and central Illinois (Wrobel *et al.*, 2008) found that many *C. albicans* strains from animals were no more similar to each other than they were to clinical strains from different continents, and concluded that there could be migration of *C. albicans* strains between humans and other animals. Phylogenetic analysis at MLST loci shows that the oak strains are no more similar to animal strains than they are to strains from humans (Supplemental Results and Figure S4; data from Wrobel *et al.*, 2008). Our findings therefore suggest that migration between humans and woodland environments is also possible.

Consistent with this conclusion, the strains isolated from oak in this study were able to grow at high temperatures (37-42°C), suggesting that they could live as mammalian commensals. Furthermore, past studies of *C. albicans* from grass and shrubs showed that environmental isolates were able to grow in rabbits and kill them within days (van Uden *et al.*, 1956; Di Menna, 1958). We do however observe more phenotypic differences among oak strains than in these early studies (van Uden *et al.*, 1956; Di Menna, 1958). More specifically, the three oak strains differ in their ability to assimilate galactose or soluble starch, to survive in 10% NaCl, or to form pseudohyphae in YM broth.

As well as divergence between strains, *C. albicans* strains from oak show high within-strain heterozygosity even after excluding LOH regions (mean 0.7%) compared to clinical strains (mean 0.6%, Table 2; Wilcoxon test, *P* = 0.003). In sexually reproducing *Saccharomyces* yeast, levels of heterozygosity also appear to differ among habitats, but in *Saccharomyces cerevisiae*, most oak strains are fully homozygous, and most non-woodland strains show average heterozygosity levels around 0.1%. Even the average heterozygosity for *S. cerevisiae* strains that were hybrids between diverged populations was lower than we see for *C. albicans* (0.4%, Table 2; Peter *et al.*, 2018). *Saccharomyces paradoxus* is similar to *S. cerevisiae* and also unlike *C. albicans* in that *S. paradoxus* oak strains are almost completely homozygous (Johnson *et al.*, 2004).

The higher heterozygosity of *C. albicans* oak strains compared to clinical *C. albicans* could arise (i) through recent mating between diverged lineages, (ii) if they represent asexual lineages that have experienced less long-term mitotic recombination than those of the average clinical strain, or (iii) because of increased natural selection for heterozygosity in the oak environment.

One of the three oak strains (NCYC 4146) belongs to clade 4 and shows no evidence for mating with another diverged clade (Figure 3). However levels of genome-wide heterozygosity for this strain were similar to those of clinical strains (Table 2) and we cannot exclude genetic exchange for the two most heterozygous strains (NCYC 4144 and NCYC 4145; Figures 3, S5 and S6). Strains from multiple clades were living within 150 meters of each other and one strain had homozygosed at the mating locus, therefore encounters between mating-capable strains from different clades are possible at cool temperatures where a parasexual cycle is most likely.

Regardless of whether higher levels of oak heterozygosity arise through parasexual or asexual cycles, there could be natural selection against deleterious alleles in homozygotes (Bougnoux *et al.*, 2008), and this selection pressure could be stronger in an open or stressful environment. The clinical strains studied here show increasing growth rates and laboratory fitness with increasing genome-wide heterozygosity but no correlated effects on virulence (Hirakawa *et al.*, 2015). Homozygous diploid strains grow poorly compared to heterozygous strains (Hickman *et al.*, 2013) and loss of heterozygosity across even small genomic regions can lead to negative fitness consequences under stress (Ciudad *et al.*, 2016). Consistent with a potential effect of increased natural selection against homozygosity, a lower proportion of *C. albicans* genomes from oak recently underwent LOH compared with clinical strains, and levels of heterozygosity were higher genome-wide in oak strains (Table 2).

### *C. albicans* lives on old oaks in an ancient wood-pasture

*C. albicans* from oak differ from clinical strains in that they are unusually heterozygous. Oak strains are highly heterozygous both because they have less DNA that recently homozygosed and because of showing heterozygosity at thousands of sites more than expected for clinical strains (Tables 2 and S1). Furthermore, the three oak strains were genetically diverged from each other (Figure 3) which implies that they do not represent laboratory contaminants from a human. Humans only rarely carry more than one distinct strain of *C. albicans*, and strains from multiple clades are especially rare (Bougnoux *et al.*, 2006). In addition, these strains were isolated from three different unusually old trees and in all cases from bark over 1.5 meters above the ground and alongside negative controls that were clear (Robinson *et al.*, 2016). The genetic divergence between oak strains also implies that these oak trees were not colonised as a result of contamination from a single animal in the woods. It is rare for domestic or wild animals to carry multiple strains of *C. albicans* and it is especially rare for these to belong to different clades (Wrobel *et al.*, 2008). These results suggest that *C. albicans* is able to live on oaks for appreciable lengths of time and therefore that it is not an obligate commensal of warm-blooded animals.

If *C. albicans* can stably inhabit a woodland environment, then why have they only been isolated from trees on a couple of occasions (Lachance *et al.*, 2011; Robinson *et al.*, 2016)? For example, three recent surveys of trees for yeast did not discover *C. albicans* but report the isolation of other *Candida* species and therefore could have detected *C. albicans* if it were present (Maganti *et al.*, 2011; Charron *et al.*, 2014; Sylvester *et al.*, 2015). All three of these surveys focused on northern North America, and it may be too cold for *C. albicans* in this region. The trees harboring *C. albicans* in the Robinson *et al.* (2016) study had larger trunk girths and were probably older than most other trees sampled in the New Forest (Table 1) and the rest of Europe. If past surveys for woodland yeast did not target old trees, it is therefore possible that *C. albicans* could have been missed.

The comparison between other human pathogenic fungi and their wild relatives on trees is proving to be important for an understanding of their pathogenicity (Gerstein and Nielsen, 2017). Futhermore, study of the natural enemies of *Cryptococcus gatii* from *C. gatii-*positive plant and soil samples could lead to the development of new antifungals (Mayer and Kronstad, 2017). If *C. albicans* is not an obligate commensal of warm-blooded animals, then comparisons between clinical and free-living strains of *C. albicans* will also be important for understanding its commensalism and pathogenicity. A major limitation in this endeavor is that very few *C. albicans* strains are available for study from non-animal sources. The few isolates that have been obtained were from a broad range of sources (van Uden *et al.*, 1956; Di Menna, 1958; Lachance *et al.*, 2011; Robinson *et al.*, 2016), and general environmental sampling for fungal pathogens can be challenging (Gerstein and Nielsen, 2017). In the future, the targeting of old trees could lead to improved environmental sampling success.

## Materials and Methods

### Yeast strains

We generated short-read genome data for the type strain of *C. albicans* (NCYC 597) and three strains of *C. albicans* from the bark of oak trees in the New Forest in the U.K (Robinson *et al.*, 2016, Table 1). Robinson *et al.* (2016) describe the methods used to isolate the oak strains. Briefly, every tree that was sampled was photographed, its trunk girth was measured in order to estimate tree age, and its longitude and latitude were recorded (Robinson *et al.*, 2016). Negative controls were generated at every field site, and these were subjected to the same procedures and handling as other samples except that no bark was inserted into negative controls after opening tubes in the field. Two of the three trees with *C. albicans* (FRI5 and FRI10) were directly associated with negative controls and there were negative controls at six of the thirty trees sampled in the New Forest (FRI5, FRI10, FRI15, OCK5, OCK10, OCK15). In total, 6 out of the 125 sample tubes collected in the New Forest (Robinson *et al.*, 2016) were negative controls. We used the reference genome sequence for *C. albicans* strain SC5314_A22 (haplotype A, version 22; GCF_000182965.3) from the NCBI reference sequence database. For comparisons of oak strain to clinical strain genomes, we used short-read genome data from the European Nucleotide Archive for 20 clinical strains (PRJNA193498; Hirakawa *et al.*, 2015), for wild-type SC5314 (SRR850113) and the related 1AA mutant (strain AF9318-1, SRR850115; Legrand *et al.*, 2008; Muzzey *et al.*, 2013). Twelve of the 20 clinical strains in the dataset from Hirakawa *et al.* (2015) were homozygous at the MTL locus, and two of these (P78042 and P75010) were homozygosed at this locus in the lab (Sahni *et al.*, 2009).

### Phenotypic profiling of the type strain and strains from oak

The type strain of *C. albicans* (NCYC 597) and the three oak strain strains were characterised biochemically, morphologically and physiologically according to the standard methods described by (Kurtzman *et al.*, 2011). The temperature for growth was determined by cultivation on YM (yeast extract-malt extract) agar. In addition, the three oak strains were subjected to the culturing and storage conditions described in (Robinson *et al.*, 2016).

### Whole-genome sequencing and base calling

Purified genomic DNA was extracted from saturated 1.5 ml cultures using a MasterPure yeast DNA purification kit (Epicentre) and following the manufacturer’s instructions. Whole genome sequencing of the four *C. albicans* genomic DNA samples was carried out at the Earlham Institute, Norwich, UK. Libraries were constructed using their LITE (Low Input Transposase Enabled) methodology for library construction of small eukaryotic genomes based on the Illumina Nextera kits. Each library pool was sequenced with a 2 × 250 bp read metric over six lanes of an Illumina HiSeq2500 sequencer. Adapters were trimmed using Trimmomatic (version 0.33, Bolger *et al.*, 2014) with default settings for paired end data and the ILLUMINACLIP tool (2:30:10). We used FastQC (version 0.11.4, http://www.bioinformatics.babraham.ac.uk/projects/fastqc/) to check read quality and the presence of adapters before and after trimming, and used trimmed, paired read data in subsequent analyses.

Short read data for all strains (3 oak strains, the type strain, 21 clinical strains and the 1AA mutant) were mapped to the SC5314_A22 reference genome using Burrows-Wheeler Aligner (bwa mem, version 0.7.10; Li and Durbin, 2009). We used SAMtools (version 1.2; Li *et al.*, 2009) to generate sorted bam files that were merged in cases where there were multiple sets of read-pair data files per strain. To generate a consensus (in genomic variant call format, gVCF) we used mpileup from SAMtools and then bcftools call (with the -c option). SAMtools mpileup was used with default settings except that maximum read depth was increased to 10,000 reads and we used the -I option so insertions and deletions were excluded.

### Verification of DNA sequence at MLST, MTL and rDNA loci

We inferred the standard genotype calls for the type strain and the three oak strains using the genomic data generated above and also verified these by independent DNA extraction, PCR and sequencing. Purified genomic DNA was extracted as above, except that 100 units of lyticase was used to degrade the fungal cell wall for each prep, prior to DNA extraction. The DNA yield from each prep was determined by fluorimetry using a Qubit 3.0 fluorometer (ThermoFisher). The seven housekeeping genes (*AAT1α*, *ACC1*, *ADP1*, *MPI1b*, *SYA1*, *VPS13* and *ZWF1b*) used routinely for *C. albicans* strain typing were PCR-amplified and sequenced following the standard protocol (Bougnoux *et al.*, 2003; Tavanti *et al.*, 2003). The *Candida albicans* MLST database (https://pubmlst.org/calbicans/) was used to determine allele identity (sequence type, ST). The mating type (*a/α*, *a/a* or *α/α*) was determined by PCR using the method described by Tavanti *et al.* (2003). The complete ITS region, encompassing ITS1, the 5.8S rRNA gene and ITS2, was PCR-amplified directly from whole yeast cell suspensions following the procedure and PCR parameters as described by James *et al.* (1996). The ITS region was amplified and sequenced using the conserved fungal primers ITS5 and ITS4 (White *et al.*, 1990).

### Estimation of ploidy and identification of loss of heterozygosity regions

We developed a perl script, vcf2allelePlot.pl (available from https://github.com/bensassonlab/scripts/) that uses R (version 3.2.3) to visualize all the base calls that differ between a strain and the reference genome (SC5314 A22). The script plots the proportion of reads that differ from the reference (the “allele ratio”, Zhu *et al.*, 2016; Todd *et al.*, 2017) at every site that has a point substitution along each chromosome (Figures 1 and S1). Following a standard approach (the B allele approach; Teo *et al.*, 2012; Yoshida *et al.*, 2013; Zhu *et al.*, 2016), we visually decided ploidy state from plots as follows. Heterozygous diploids were called where chromosomes had allele ratios of 1 and 0.5. Heterozygous triploids were called where chromosomes had allele ratios of 0.33, 0.66, and 1.

The same script used to visualize allele ratios (vcf2allelePlot.pl) was used to identify LOH regions as follows. The genome was divided into 100 kb non-overlapping windows and LOH was called for windows where the proportion of heterozygous sites was below 0.1%. We estimated the proportion of heterozygous sites after excluding low quality sites (phred-scaled consensus quality below 40), centromeres, annotated repeats and sites in each window with over double the average genome-wide read depth for a strain. Sites were considered heterozygous if their allele ratio was between 0.2 and 0.8. In a separate analysis, we used histograms generated in R to decide the 0.1% threshold for the identification of LOH regions (Figures 2 and S3).

There were cases where strains were predominantly diploid, but had allele ratios of 1 all along one or more chromosomes, and this could result from monosomy or from whole-chromosome LOH. In order to distinguish between these two possibilities, a second approach is often used to test for aneuploidy (Teo *et al.*, 2012; Zhu *et al.*, 2016), and we also tested that approach here. In cases of aneuploidy, read depths will differ between chromosomes. We used SAMtools depth (version 1.3.1; Li *et al.*, 2009) with maximum read depth (the -d option) set to 10,000 to estimate read depth for each position, then R was used for sliding window statistical analyses and visualization. We estimated average read depth for non-overlapping 1 kb windows across each chromosome, and then estimated a median from these for each chromosome. Chromosomes were then considered aneuploid if median read depths in pairwise comparisons between chromosomes differed by over 35% (Zhu *et al.*, 2016). However, this approach assumes random fragmentation of DNA prior to sequencing and therefore even read depth across genomic regions. While this assumption holds relatively well when DNA is mechanically sheared, enzymatic approaches for DNA fragmentation are more likely to result in uneven read depth that correlates with base composition (Marine *et al.*, 2011; Quail *et al.*, 2012; Teo *et al.*, 2012). The genome data that was generated as part of this study for the type and oak strains was generated using an enzymatic fragmentation protocol and showed continuously uneven read depth within and between chromosomes (Figure S2). Therefore, we were unable to use the read depth approach to test for aneuploidy. As a result, we rely on the base calling approach which is best for determining overall ploidy, but cannot detect aneuploidy for chromosomes that are homozygous (chromosomes 5 and 7 of strain NCYC 4144 from oak, Figure 1).

### Phylogenetic analysis and *in silico* chromosome painting

In order to determine the relationships between strains, we used a maximum likelihood phylogenetic approach implemented in RAxML (version 8.1.20, Stamatakis, 2014). Using seqtk from SAMtools, we converted base calls in gVCF format to fasta format sequence and filtered bases that had quality scores below a phred-scaled quality score of 40 (equivalent to an error rate of 1 in 10,000). All genome sequences were mapped against the reference genome, and were therefore already aligned against it because insertions and deletions were excluded. Fasta format alignment files were converted to phylip format using fa2phylip.pl (https://github.com/bensassonlab/scripts/). For all phylogenetic analyses, we used RAxML with a general time reversible evolutionary model and a γ distribution to estimate heterogeneity in base substitution rates among sites (GTRGAMMA), and 1,000 bootstrap replicates. For genome-wide phylogenetic analysis, we included genome data for all strains including the reference genome, and concatenated the alignments for every chromosome into a single genome-wide alignment. For phylogenetic analysis of short genomic regions within chromosomes, we extracted alignments for phylogenetic analysis using faChooseSubseq.pl (https://github.com/bensassonlab/scripts/).

In order to test whether a strain is similar to a single clade in all parts of its genome, we developed faChrompaint.pl (https://github.com/bensassonlab/scripts/) to “paint” chromosomes according to similarity to known clades. Several other tools already exist for painting chromosomes *in silico* in order to identify admixture between populations (reviewed in Schraiber and Akey 2015), however these require phased haplotype data which are not available for *C. albicans*. The <u>faChrompaint.pl</u> script takes fasta formatted whole-chromosome alignments as input, and divides the genome of a study strain into non-overlapping windows (we set the window size to 100 kb). The script uses R to generate a plot with every window colored according to the clade assignment of the most similar strain in that window. The most similar DNA sequence was the one with the lowest proportion of differing sites. We used the clade assignments made by Hirakawa *et al.* (2015) for their 21 clinical strain genomes to define clades, and colored windows green if their greatest similarity was to an oak strain sequence. If a strain is genetically diverged from the seven clades studied by Hirakawa *et al.* (2015), then similarity to a known clade does not necessarily imply recent common ancestry. We therefore filtered diverged regions by leaving those windows blank. More specifically, we did not color windows with over 0.066% divergence from known clades because most within-clade pairwise comparisons (90%) show divergence levels below 0.066%, while most between-clade comparisons show divergence above 0.066% (Figure S8).

We also tested whether similarities to different clades resulted in statistically supported phylogenetic incongruence in homozygous regions. For two oak strains (NCYC 4144 and NCYC 4145), most homozygous regions showed high divergence from known clades. We therefore ran <u>faChrompaint.pl</u> without applying a divergence filter, and identified regions likely to show incongruent phylogenies, then compared phylogenetic analyses between these regions. This chromosome painting approach was successful in identifying regions with phylogenetic incongruence (see examples in Figures S5 and S6).

### Estimation of levels of heterozygosity

In order to estimate levels of heterozygosity either genome-wide (Table 2) or in 100 kb non-overlapping windows (Figure 2), we estimated the proportion of sites that were heterozygous. For all estimates of levels of heterozygosity, only high quality sites (phred-scaled quality over 40) were considered. Sites were considered heterozygous if the proportion of sites differing from the reference sequence (the allele ratio) was between 0.2 and 0.8. In a diploid, it is also possible for sites to be heterozygous with an allele ratio of 1 in cases where 3 alleles exist for a site because both alleles could differ from that of the reference genome. For example, the reference genome may have an A at a site, and a study strain could show an allele ratio of 1 while being heterozygous for C and T alleles. However, levels of intraspecific genetic diversity are sufficiently low that we expect triallelic sites to represent a small proportion of all heterozygous sites, and therefore not to affect our conclusions. For example, if the true proportion of heterozygous sites is 0.007 (close to the levels we observe in Table 2), then the expected proportion of sites with a second point substitution would be 4.9 × 10^−5^ (i.e. 0.007^2^). The observed number of high quality triallelic sites in each (14 Mbp) genome sequence are slightly lower than expected: up to 1 × 10^−5^ (144 sites) for the oak strains and all clinical strains except the type strain. The type strain (NCYC 597), which is mostly triploid, has the largest number of triallelic sites (173 sites). Differences between oak and clinical strains in the exclusion of these few sites cannot explain the higher levels of heterozygosity seen for oak strains which exceed that of clinical strains by thousands of sites (Table 2).

## Data Accessibility

DNA sequences determined for this study are available in EBI’s ENA as PRxxx. Perl scripts are available at https://github.com/bensassonlab/scripts. The type strain and *C. albicans* strains isolated from oak are available from the National Collection of Yeast Cultures in the U.K..

## Acknowledgements

We thank Darren Heavens of the Earlham Institute, Norwich, UK for the LITE-based whole genome sequencing of the oak and type strains of *C. albicans*. We also thank Casey Bergman, Dave Hall and Jenna Hamlin for helpful comments on the manuscript. The work of J.D., C.J.B., A.E. and I.N.R was supported by the Biotechnology and Biological Sciences Research Council, through a National Capability award to the National Collection of Yeast Cultures (grant number BBS/E/F/00044440 to I.N.R.) and an Institute Strategic Programme award to the Institute of Food Research (grant number BBS/E/F/00044471 to I.N.R.).

## Authors’ Contributions

D.B., J.D. I.N.R. and S.A.J. conceived and designed the research; S.A.J., C.J.B. and A.E. generated the data; D.B., J.D., J.M.L. and S.A.J. analyzed the data; and D.B. wrote the manuscript with contributions from J.D., S.A.J. and I.N.R.

## Supplemental Files

1. Bensasson_etalTableS1.tsv: a table in text format with tab separated values summarizing heterozygosity analyses for every strain. This includes exact counts of high quality heterozygous base calls (highQualityHetCount); the total length of high quality sequence (highQualityLength; bases with a phred-scaled quality score over 40); the proportion of high quality heterozygous sites; the length of regions that have undergone Loss of Heterozygosity (LOHlength) assessed in 100 kb windows; heterozygosity analysis after excluding LOH regions, centromeres and annotated repeats (annotationLohFilteredHetCount, annotationLohFilteredLength, annotationLohFilteredHeterozygosity); heterozygosity analysis after excluding LOH regions, centromeres and annotated repeats, and regions with more than double the expected read depth (depthFilteredHetCount, depthFilteredLength, depthFilteredHeterozygosity); heterozygosity analysis at 948,860 nucleotide sites that are common to all strains (sitesIn950kbHetCount, sitesIn950kbLength, sitesIn950kbHeterozygosity).
2. Bensasson etalSupp.pdf: a pdf file with Supplemental Results and Figures S1 to S8. Supplemental Results describe a MLST analysis that shows oak strains are as similar to clinical strains as they are to *C. albicans* from animals.

### Summary of Supplemental Figures in Bensasson etalSupp.pdf

1. Figure S1 showing the base calling plots used to estimate ploidy and to visualize LOH regions. Ploidy was estimated based on the proportion of base calls differing from the reference at every site in the genome where there is a nucleotide substitution for each clinical strain. This analysis confirms 6 cases of aneuploidy identified by Hirakawa et al (2015): 12C chr4, 19F chr7, L26 chr7, P60002 chr4 and chr6, P78042 chr4.
2. Figure S2 showing read depth across the genome of each strain estimated in 1 kb non-overlapping sliding windows. Read depth was continuously uneven within and between chromosomes for the type strain (NCYC 597) and oak strains (NCYC 4144, NCY 4145, NCYC 4146). This is a problem if a ploidy estimation approach assumes discrete jumps in read depth between chromosomes. In contrast, the assumption of discrete jumps in read depth between chromosomes holds much better for the analysis of the data generated by Hirakawa *et al.* (2015), and our estimates confirm all their aneuploidy calls.
3. Figure S3 showing the distribution of levels of heterozygosity estimated in 100 kb non-overlapping windows across the genome of each strain.
4. Figure S4 showing a. phylogenetic relationships between clinical strains and oak strains using only data from MLST loci. b. phylogenetic relationships between clinical strains, oak strains and animal strains. Oak strains (purple) are more similar to clinical strains than animal strains, which are prefixed with “ST”. Sequence types and clade assignments for domestic and wild animals were determined by Wrobel *et al.* (2008) and sequences were downloaded from http://pubmlst.org/calbicans/.
5. Figure S5 showing chromosome-by-chromosome maximum likelihood trees for clinical strains and oak strains.
6. Figure S6 showing that one oak strain (NCYC 4144) shows phylogenetic incongruence in different parts of the genome.
7. Figure S7 showing the genomes of each clinical strain, split into 100 kb windows and colored according to the clade assignment of the most similar clinical strain. In cases where the level of similarity is above that expected for 90% of within-clade comparisons, the 100 kb window is coloured white. a. clade 1 strains; b. clade 2 strains; c. clade 3 strains; d. clade 4 strains; e. clade 6, 8 and 11 strains.
8. Figure S8 showing histograms used to visualize within-clade divergences in 100 kb windows (blue), and to compare these to between-clade divergences (purple). Most within-clade divergences (90%) are below 0.066% (green line) while most between clade divergences are above it. In cases where sequence divergence between sequences is above a threshold of 0.066% chromosomes were painted white in Figures 3b and S7 to show that they were too diverged from other sequences for clade assignment.

